# DiscoMark: Nuclear marker discovery from orthologous sequences using draft genome data

**DOI:** 10.1101/047282

**Authors:** Sereina Rutschmann, Harald Detering, Sabrina Simon, Jakob Fredslund, Michael T. Monaghan

## Abstract

High-throughput sequencing has laid the foundation for fast and cost-effective development of phylogenetic markers. Here we present the program DISCOMARK, which streamlines the development of nuclear DNA (nDNA) markers from whole-genome (or whole-transcriptome) sequencing data, combining local alignment, alignment trimming, reference mapping and primer design based on multiple sequence alignments in order to design primer pairs from input orthologous sequences. In order to demonstrate the suitability of DISCOMARK we designed markers for two groups of species, one consisting of closely related species and one group of distantly related species. For the closely related members of the species complex of *Cloeon dipterum* s.l. (Insecta, Ephemeroptera), the program discovered a total of 78 markers. Among these, we selected eight markers for amplification and Sanger sequencing. The exon sequence alignments (2,526 base pairs (bp)) were used to reconstruct a well supported phylogeny and to infer clearly structured haplotype networks. For the distantly related species we designed primers for several families in the insect order Ephemeroptera, using available genomic data from four sequenced species. We developed primer pairs for 23 markers that are designed to amplify across several families. The DISCOMARK program will enhance the development of new nDNA markers by providing a streamlined, automated approach to perform genome-scale scans for phylogenetic markers. The program is written in Python, released under a public license (GNU GPL v2), and together with a manual and example data set available at: https://github.com/hdetering/discomark.

## Introduction

The inference of phylogenetic relationships has benefited profoundly from the availability of nuclear DNA (nDNA) sequences for an increasing number of organism groups. The development of new phylogenetic markers has provided unprecedented insight into the evolutionary relationships of non-model organisms in particular (Ellegren 2014). Large sets of nDNA markers (single copy genes) have recently been designed for taxonomic groups for which genomic resources were available, e.g. cichlid fish (Meyer *et al.* 2015), ray-finned fish (Near *et al.* 2012), reptiles (Ruane *et al.* 2014), birds (Kerr *et al.* 2014) and flowering plants (Zeng *et al.* 2014). However, for many taxonomic groups there are only a handful of nDNA markers available that are suitable for phylogenetic reconstruction. Other approaches, such as ultra-conserved element (UCE) sequencing (Faircloth *et al.* 2012), anchored hybrid enrichment (Lemmon and Lemmon 2012), restriction site-associated DNA (RAD) sequencing (Baird *et al.* 2008) or genotyping by sequencing (GBS, Elshire *et al.* 2011) have become popular for addressing specific questions in systematics or population genetics; however, these methods are still cost-intensive, require a comparatively high amount of starting DNA material and can depend on the availability of reference genomes (e.g. anchored hybrid enrichment). Consequently, standard Sanger sequencing approaches are still in high demand for various research questions.

Identification of novel phylogenetic markers has been a predominantly manual process, which impedes their large-scale development, and comprehensive primer design based on large sets of multiple sequence alignments remains challenging. Existing tools can generally be classified into those developed for primer design or for marker discovery. Primer design tools can design primer pairs for single loci (e.g. GEMI, Sobhy *et al.* 2012; PRIMER3, Untergasser *et al.* 2012; CEMASUITE, Lane *et al.* 2015) or multiple loci at once (BATCHPRIMER3, You *et al.* 2008; PRIMERVIEW, O’Halloran 2015). Some programs are specific for highly variable DNA targets (PRIMERDESIGN, Brodin *et al.* 2013; PRIMERDESIGN-M, Yoon and Leitner 2015), viral genomes (PRISM, Yu *et al.* 2015), and transcriptome input data (SCRIMER, Morkovsky *et al.* 2015). Marker discovery tools target single nucleotide polymorphism (SNP) markers in polyploid organisms (POLYMARKER, Ramirez-Gonzalez *et al.* 2015), and putative single copy loci from plant transcriptomes (MARKERMINER, Chamala *et al.* 2015). The challenge of developing new phylogenetic markers lies both in the discovery of conserved regions, the design of primer pairs and an estimation of the level of their phylogenetic signal.

Here we present DISCOMARK (=Discovery of Markers), a flexible, user-friendly program that identifies conserved regions and designs primers based on multiple sequence alignments taken from FASTA-formatted files of putative orthologous sequences from whole-genome or whole-transcriptome data. The program can be used to easily screen for phylogenetically suitable nDNA markers and to design primers that can be used for Sanger sequencing as well as high-throughput sequencing. The program is structured into several steps that can be individually optimized by the user and run independently. In terms of input, the program can be applied on large and small sets of taxa, including both closely and distantly related species. Ideally, orthologous sequences in combination with a whole-genome reference sequence are used. Thus, exon/intron boundaries can be inferred using the reference for each marker. Under the default settings, the program will design several primer pairs that anneal in conserved regions. The visualization of the alignments with potential primers allows the user to choose between primers targeting exons or introns (e.g. exon-primed intron-crossing (EPIC) markers). Additionally, information about the suitability as phylogenetic markers is provided by an estimate of the number of SNPs per marker and the applicability across species. Finally, we demonstrate the utility of DISCOMARK for (1) closely related species (i.e. *Cloeon dipterum* s.l. species complex) using whole-genome data, and (2) distantly related species (i.e. insect order Ephemeroptera) using whole-genome data derived from genome sequencing projects. in order to generate genomic reference sequences we used draft whole genome sequencing at shallow coverage followed by draft genome assembly. In one scenario (*C. dipterum* s.l. species complex) we demonstrate that incomplete genomic data can be used for ortholog prediction and primer design as well.

## Materials and Methods

### DISCOMARK implementation

The program DISCOMARK is written in Python and was developed to design primer pairs in conserved regions of predicted orthologous genes. orthologs are required for phylogenetic studies. The ortholog identification step is not part of the DISCOMARK workflow but DISCOMARK is designed to directly work with the output of several ortholog prediction programs, e.g. HAMSTR (Ebersberger *et al.* 2009), or Orthograph (https://github.com/mptrsen/Orthograph, last accessed March 25, 2016). Orthologous groups may be derived from genome or transcriptome sequence data. In addition to the orthologous genes, genomic data such as whole-genome sequencing data can be provided to DISCOMARK as a guide to detect exon/intron boundaries. DISCOMARK performs seven steps, combining Python scripts with widely used bioinformatics programs (Fig. 1). The steps: (1) combine orthologous groups of sequences, (2) align sequences of each orthologous group using MAFFT v.7.205 (Katoh and Standley 2013), (3) trim sequence alignments with TRIMAL v.1.4 (Capella-Gutierrez *et al.* 2009), (4) align sequences against a reference (e.g. whole-genome dataset from the same or closely related taxa) with BLASTN v.2.2.29 (Altschul *et al.* 1997; Camacho *et al.* 2009) and re-alignment using MAFFT, (5) design primer pairs on single-gene alignments using a modified version of PRIFI (Fredslund *et al.* 2005), adapted by us into a Python package that uses BioPython v.1.65 and Python v.3.4.3, (6) check primer specificity with BLASTN, and (7) generate output in several formats (visual HTML report, tabular data and FASTA files of the primers). The results of each step can be inspected in the respective output folders.

**Fig. 1.**
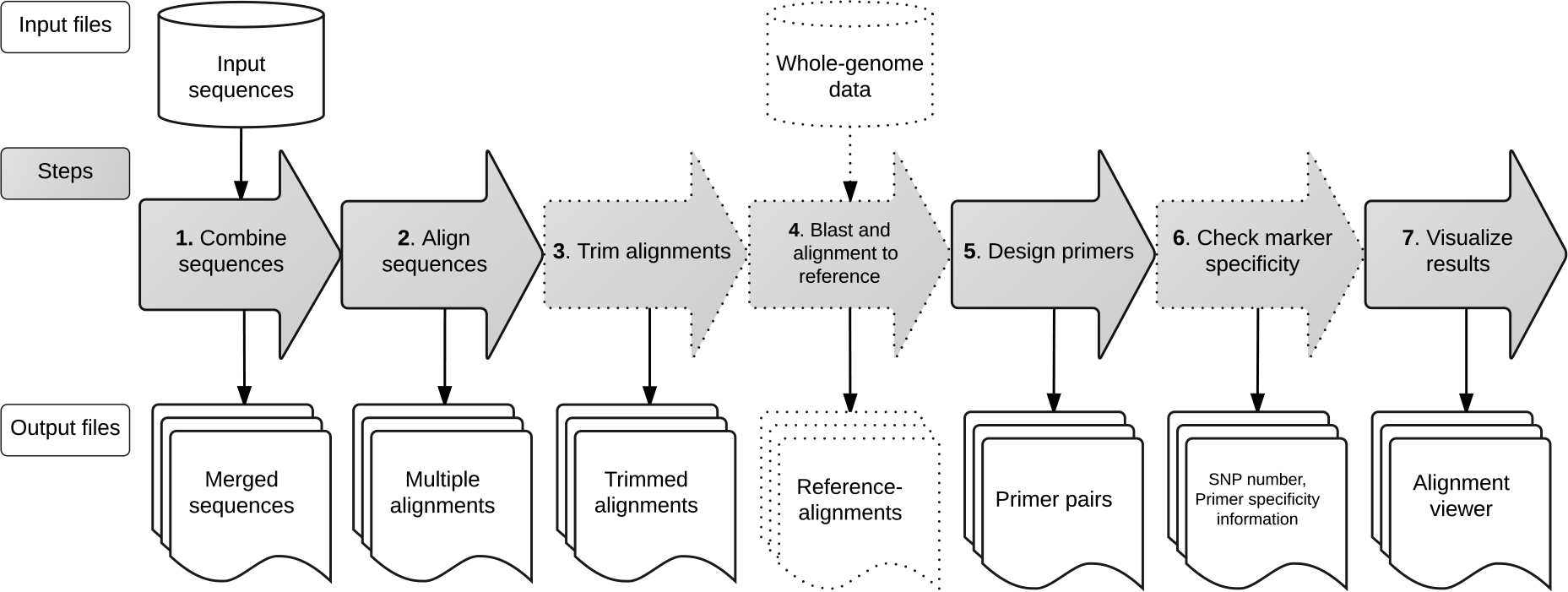
Overview of the DISCOMARK workflow and processing steps. Arrows with a broken outline indicate optional steps (for details see Materials and Methods section).

1. *Combine sequences*. In the first step, the putative orthologous sequences of different taxa are combined according to the orthologous groups. The input files are expected to be nucleotide sequences in FASTA format. We recommend using putative orthologous exon sequences (e.g. CDS) in combination with whole-genome data (e.g. a draft genome assembly). Each input file is expected to contain the sequences of one orthologous group; orthologs of each input taxon are to be organized into a taxon folder. Importantly, file names represent the ortholog identifiers used to combine orthologous sequences of the various input taxa; by default, ortholog prediction tools follow that convention.
2. *Align sequences*. Orthologous sequences combined according to the orthologous groups are separately aligned with the multiple sequence alignment (MSA) program MAFFT. Alignment parameters can be specified by the user via a configuration file (DISCOMARK.conf, located in the program folder). Default parameters are the following: ‘--localpair --maxiterate 16 -- inputorder --preservecase --quiet’ (L-INS-i alignment method). We chose MAFFT as multiple alignment tool because it combines accuracy and efficiency and has been adopted widely in the scientific community (Pais *et al.* 2014; Szitenberg *et al.* 2015).
3. *Trim alignments*. In order to remove poorly aligned regions, sequence alignments are trimmed using TRIMAL. The program TRIMAL analyzes the distribution of gaps and mismatches in the alignment and discard alignment positions and sequences of low quality. By default, DISCOMARK calls TRIMAL with the ‘-strictplus’ method. The preset is used by TRIMAL to derive the specific thresholds for alignment trimming (minimum gap score, minimum residue similarity score, conserved block size). Since alignment trimming largely depends on the input data and influences the downstream results, TRIMAL can also be run with different settings (e.g. ‘-gappyout’, ‘-strict’, ‘-automated1’ but see Capella-Gutierrez *et al.* 2009. Alternatively, there is also the option to deactivate the alignment trimming with the DISCOMARK option ‘--no-trim’ or use alternative trimming programs such as GBLOCKS (Castresana 2000; Talavera and Castresana 2007) or GUIDANCE2 (Landan and Graur 2008; Sela *et al.* 2015).
4. *Blast and alignment to reference*. In this step a genomic reference sequence for each input ortholog is identified and added to the trimmed alignment. This step is particularly important when working with coding sequences which do not contain intron sequences; thus, a genomic sequence is needed to infer intron/exon boundaries. Working with coding sequences is advisable for more distantly related taxa which may include intron length polymorphisms, or to target EPIC markers. Any whole-genome data set (from one of the included taxa or a closely related taxa) can be used as reference for mapping the ortholog sequences. Here, mapping means that the input sequences are compared to the reference sequences, which are defined by the user using the local alignment program BLASTN. The best locally aligning reference sequence (the one that yields the longest alignment among all input sequences) for each orthologous group is added to the corresponding sequence alignment. Reference sequences are cut to 100 base pairs (bp) upstream and downstream of the first, respectively last, BLAST hit to avoid alignment length inflation. Then, the extended alignments are realigned with MAFFT. Finally, the best BLAST hits of all sequences belonging to a marker are compared and a ‘uniq_ref’ flag is set if they all map to the same reference sequence. The reference alignment step is optional; however, the inclusion of whole-genome data is essential for estimating intron/exon boundaries. Given that information, the focus of target sequences to be amplified can be on entire exon markers, EPIC markers, or a combination.
5. *Design primers*. The single-gene alignments, after trimming, mapping and re-aligning to a reference, are used as input to design primer pairs. We integrated the webtool PRIFI(http://cgi-www.daimi.au.dk/cgi-chili/PriFi/main, last accessed December 20, 2015) as a Python package that provides a comprehensive set of parameters. As default settings for DISCOMARK we chose the following: estimated product length between 200-1,000 bp (‘OptimalProductLength = [400,600,800,1000], MinProductLength = 200, MaxProductLength = 1000’), maximum number of ambiguity positions within the primer sequences (‘MaxMismatches = 2’), primer length between 20-30 bp (‘MinPrimerLength = 20, MaxPrimerLength = 30, OptimalPrimerLength = [20, 25]’), melting temperature of the primer pairs between 50-60°C (‘MinTm = 50.0, MinTmWithMismatchesAllowed = 58.0, SuggestedMaxTm = 60.0’), and we set the maximum number of primer pairs per alignment to six (note: only settings different from the PRIFIdefault are mentioned above). The program PRIFIwas originally developed to design intron-spanning markers (but see Fredslund et al. 2005). Here we use it because it enables primer design based on MSA input. Parameters for PRIFIcan be specified in the DISCOMARK configuration file (‘discomark.conf’).
6. *Check marker specificity*. To ensure the specificity of the designed primer pairs, we compare their sequences against the NCBI database (‘refseq_mrna’). Primer sequences are searched in the NCBI database (‘refseq_mrna’) using the online BLASTN interface. The default search settings are restricted to human and bacterial targets using the Entrez query ‘txid2[ORGN] OR txid9606[ORGN]’ because these are most likely to be present as contaminants in sequencing libraries. The result hits of the BLAST search are indicated to the user in the HTML output.
7. *Visualize results*. As final step, the program produces a HTML report containing the list of designed primers, an alignment viewer and plots visualizing the discovered set of markers (Fig. 2). Besides the primer sequences the report lists several features such as the melting temperatures, predicted sequence length, and the number of taxa amplified by each primer set (Fig. 2a). Selected primer pairs and primer lists can be downloaded as FASTA or CSV files, respectively. The report highlights the species coverage achieved by each discovered marker, i.e. how many species’ sequences each primer set is expected to amplify, as an estimate of how universally each primer set can be applied. An identification of uniqueness within the reference genome (as an estimator of single-copy status) is given within the flag ‘uref’, starting whether all sequences belonging to a marker had their best BLAST hit to the same reference sequence. Additionally, functional annotations are reported, if available, to guide the user in the selection of markers of interest. Annotations can be supplied in form of a tab-delimited file with the ‘-a’ option. In principle, any kind of annotations can be used depending on the desired research objective. In our usage scenarios, we used gene ontology (GO) terms which were retrieved by mapping the gene IDs contained in the HAMSTR core ortholog set via the UniProt website (http://www.uniprot.org/, last accessed December 20, 2015). common input ortholog sequences are visualized (Fig. 2b). in order to provide a measure of the suitability of the markers for phylogenetic reconstruction the program calculates the number of SNPs between a primer pair by comparing the aligned input sequences against each other. The number of SNPs between each primer pair is visualized in relation to the estimated product length (Fig. 2c) and reported in the tabular output.

**Fig. 2.**
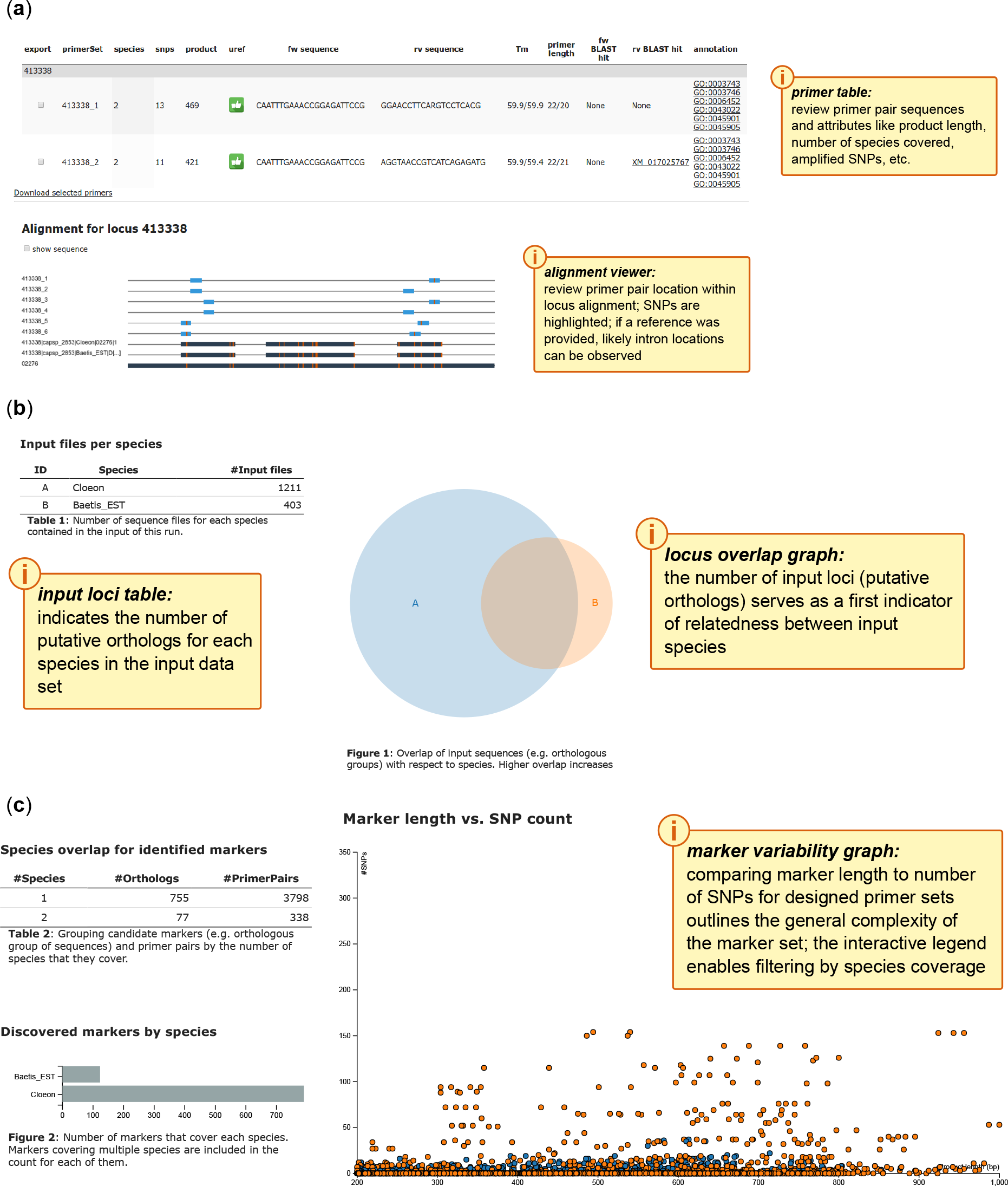
Schematic visualization of DISCOMARK HTML output, (**a**), overview of designed primers and view of alignment, including suggested primer pairs, (**b**), overview of provided input files, including a graph with common putative orthologous sequences, (**c**) information about the output, including identified markers, discovered markers per species, and scatter plot displaying the number of single nucleotide polymorphisms (SNPs) versus product length for each primer pair

### Usage cases

*Closely related species - Cloeon dipterum s.l. species complex*. To test the suitability of DISCOMARK for closely related species, we designed primer pairs for the species complex of *C. dipterum* s.l. (Ephemeroptera: Baetidae). The species complex consists of several closely related species, including *Cloeon peregrinator* GATTOLLIAT & SARTORI, 2008 from Madeira (Gattolliat *et al.* 2008; Rutschmann *et al.* 2014). As input to design the primer pairs, we used whole-genome sequencing data of *C. dipterum* L. 1761 and expressed sequence tags (EST) of *Baetis* sp. (Table 1). The sequence reads of *C. dipterum* were trimmed and *de novo* assembled using NEWBLER v.2.5.3 (454 Life Science Corporation) under the default settings for large datasets. Ortholog sequences prediction of both data sets was performed with HaMStR v.9 using the insecta_hmmer3-2 core reference taxa set (http://www.deep-phylogeny.org/hamstr/download/datasets/hmmer3/insectahmmer3-2.tar.gz, last accessed December 20, 2015), including 1,579 orthologous genes. We ran the program DISCOMARK with default settings (‘python run_project.py -i input/Cloeon -i input/Baetis -r input/reference/Cloeon.fa -a input/co2go.ixosc.csv -d output/cloeon_baetis’), using the predicted orthologs from HAMSTR and the whole-genome *Cloeon*-data as reference (step 4). For comparison, we also ran DISCOMARK without a reference and also present these results. The Pearson correlation between the number of SNPs located between primer pairs and corresponding estimated product length was calculated using the function cor within the stats package for R (R Development Core Team, 2016). A t-test for significance was performed using the function cor.test.

**Table 1.**
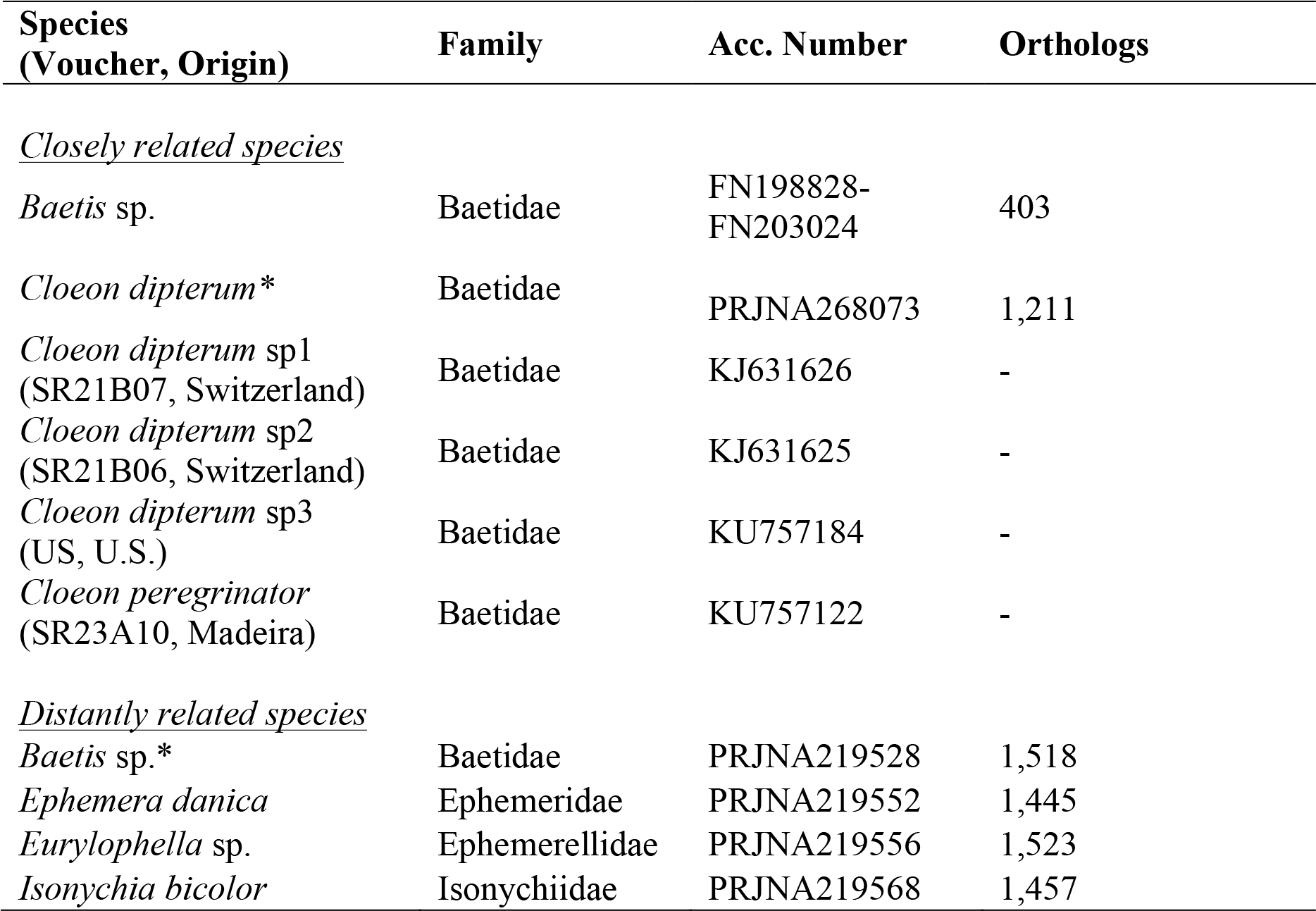
List of species used for the usage examples of the closely related species (*Cloeon dipterum* s.l. species complex), and the distantly related species (insect order Ephemeroptera).Voucher number, geographic origin, and GenBank accession numbers of cytochrome *c* oxidase subunit 1 (*cox1*) gene are given for the specimens used for the marker amplification. GenBank accession numbers of the genome data and the number of putative orthologs are given for the species used to run DISCOMARK. For both data sets species used as reference are indicated with *.

From the total of designed primer pairs (78 markers, 445 primer pairs, see results) we selected eight and amplified them for four species of the *C. dipterum* species complex (Table 1) in the laboratory. The eight markers were manually selected based on the following criteria: best alignment (i.e. EPIC markers), longest product length, and most species covered. We used standardized polymerase chain reactions (PCR; 35-40 PCR cycles with annealing temperature of 55°C), followed by Sanger sequencing. Forward and reverse sequences were assembled and edited with GENEIOUS R7 v.7.1.3 (Biomatters Ltd.), indicating ambiguous positions following the IUPAC nucleotide codes. Sequences containing heterozygous indels (i.e. alleles with length polymorphisms) were phased with CODONCODEALIGNER v.3.5.6 (CodonCode Corporation). For this we used the implemented ‘Split Heterozygous Indels’ function. Multiple sequence alignments were created for all sequences for each marker. The predicted orthologous sequences of *Baetis* sp. were used as reference to infer the exon-intron splicing boundaries (canonical and non-canonical splice site pairs). The final sequence alignments were checked for the occurrence of stop codons and indels, and split into exon and intron parts using a custom Python script (https://github.com/srutschmann/pythonscripts, last accessed March 28, 2016). Sequence alignments were phased using the program PHASE v.2.1.1 (Stephens *et al.* 2001; Stephens and Donnelly 2003) with a cutoff value of 0.6 (Harrigan *et al.* 2008; Garrick *et al.* 2010), whereby input and output files were formatted using the Perl scripts included in SEQPHASE (Flot 2010). Heterozygous sites that could not be resolved were coded as ambiguity codes for subsequent analyses. After phasing, all alignments were re-aligned with MAFFT. The number of variable and informative sites, and the nucleotide diversity per exon alignment was calculated with a custom script.

To investigate the heterogeneity of each marker’s DNA sequences, we reconstructed haplotype networks, using FITCHI (Matschiner 2016). As input for each marker we inferred a gene tree using the program RAxML v.8 (Stamatakis 2014) with the GTRCAT model and 1,000 bootstrap replicates under the rapid bootstrap algorithm. The phylogenetic relationships were calculated with Bayesian inference, using MRBAYES v.3.2.3 (Ronquist *et al.* 2012) based on a concatenated nDNA matrix that consisted of the exon sequences from all 15 nDNA markers. The best-fitting model of molecular evolution for each sequence alignment was selected via a BIC criterion in JMODELTEST v.2.1 (Guindon and Gascuel 2003; Darriba *et al.* 2012). We calculated 10^6^ generations with random seed, a burn-in of 25% and four MCMC chains. As an outgroup we used the predicted orthologous sequences of *Baetis* sp․.

*Distantly related species - insect order Ephemeroptera*. In this test case, we used contigs derived from whole-genome sequencing projects of the species *Baetis* sp., *Ephemera danica* MÜLLER 1764, *Eurylophella* sp., and *Isonychia bicolor* Walker 1853 (Table 1). The contigs from each species were used for ortholog predicting with HAMSTR v.13.2.4 (http://sourceforge.net/projects/hamstr/files/hamstr.v13.2.4.tar.gz, last accessed December 20, 2015). We ran DISCOMARK with the default settings twice. For the first run, we used the *Baetis* sp. data as reference (‘python run_project.py -i input/Baetis -i input/Ephemera -i input/Eurylophella -i input/Isonychia -r input/reference/Baetis.fa -a input/co2go.ixosc.csv -d output/mayflies’). The second run was performed without a reference (‘python run_project.py -i input/Baetis -i input/Ephemera -i input/Eurylophella -i input/Isonychia -a input/co2go.ixosc.csv -d output/mayflies_without_reference’).

## Results

### Closely related species - species complex of Cloeon dipterum s.l

Using a reference, DISCOMARK identified 78 nDNA markers with 445 primer pairs for orthologous sequences of both species (*Baetis* sp. and *C. dipterum* s.l.). Ortholog prediction yielded 403 orthologous sequences for the *Baetis* sp. EST-data and 1,211 for *C. dipterum*. For the individual species, DISCOMARK identified 793 markers for *C. dipterum* and 123 for *Baetis* sp. Markers including both species were between 200 and 931 bp with median length of 412 bp long. Their number of SNPs per marker ranged from zero to 82 (median: five) with an average of one SNP per 68 bp. Marker length and number of SNPs of the common markers were correlated with a Pearson's correlation coefficient of 0.28 (Pearson's product-moment correlation *P* < 0.001). Without using a reference, DISCOMARK identified 73 markers with 460 primer pairs for orthologous sequences of both species. The total run time for this data set on a local Linux machine (quad-core Intel i5, 8 GB RAM) was < 30 min.

The haplotype networks based on the eight selected markers showed a clear structure for all markers, including two markers with shared haplotypes for the two species from the U.S. and Madeira (Fig. 3 and Fig. S1, Supporting information). The length of the concatenated sequence alignment of the eight markers was 3,530 bp (2,526 bp exon sequence, Table S1, Supporting information). The exon sequence matrix contained 78 variable sites, 27 informative sites, and was 92.6% complete. The nucleotide diversity ranged between 0.009 and 0.028 (median: 0.013). Phylogenetic tree reconstruction based on these eight markers resulted in a phylogeny with fully resolved nodes (Bayesian posterior probability (PP) ≥ 95% Fig. 3a).

**Fig. 3.**
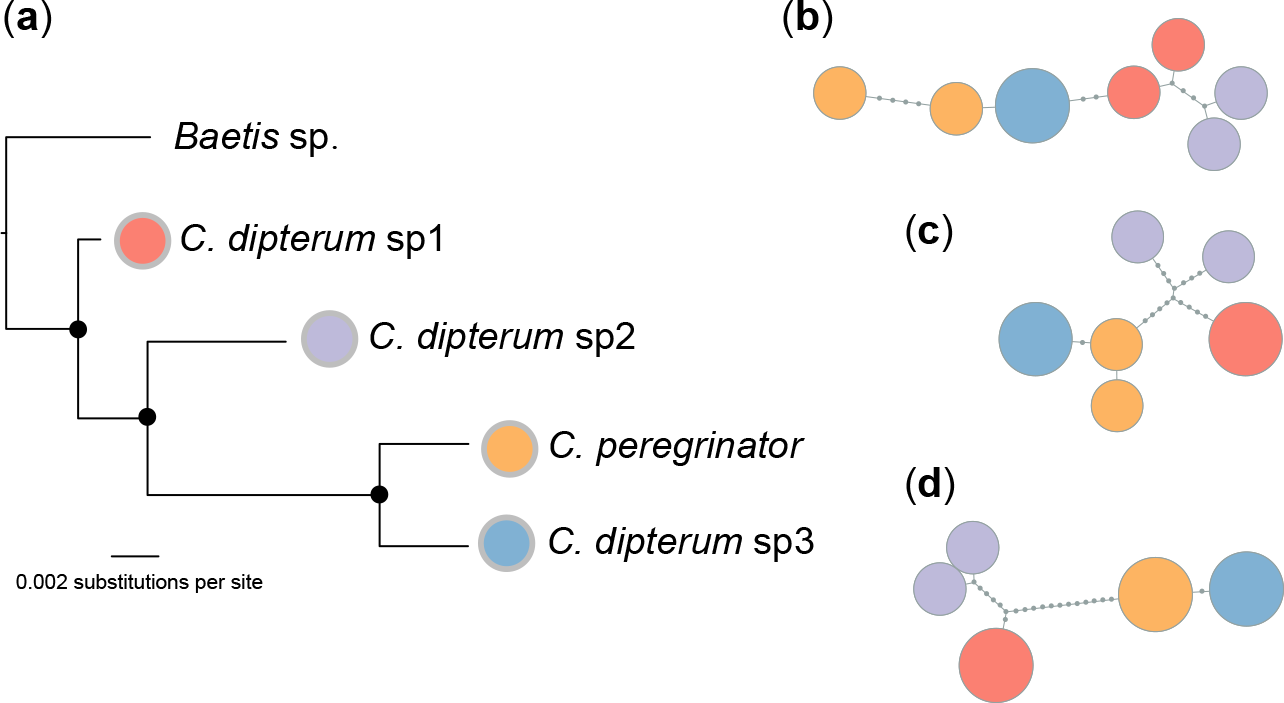
Phylogenetic reconstruction and haplotype networks for the empirical data. (**a**),Phylogenetic reconstruction of four representatives of the species complex *Cloeon dipterum* s.l., including *C. peregrinator*, based on the exon sequences of the eight newly developed nuclear DNA markers (2,526 base pairs). Bayesian inference was used to reconstruct the tree based on the concatenated supermatrix alignment. Bayesian posterior probabilities ≥ 95% are indicated by filled circles. *Baetis* was used as an outgroup. Scale bar represents substitutions per site. (**b-d**), Haplotype networks of three amplified markers, (**b**), marker 412045, (**c**), marker 412741, (**d**), marker 412048 (full set of haplotype networks is available in Fig. S1, Supporting information). Circles are proportional to haplotype frequencies. Small circles along the branch indicate missing or unsampled haplotypes. Colors correspond to the four putative species.

The species *C. dipterum* sp1 was found as outgroup to a clade containing the species *C. dipterum* sp2 from Switzerland and the two species from the U.S and Madeira. The latter two species formed a monophyletic clade. The use of the marker set for *C. dipterum* developed here resulted in a fully resolved phylogenetic tree in contrast to the mitochondrial tree of Rutschmann *et al.* (2014). The phylogeny of the *Cloeon*-species complex presented here also is in complete agreement with species tree reconstructions based on 59 nuclear DNA markers (Rutschmann *et al.,* submitted).

### Distantly related species - insect order Ephemeroptera

In total, we found 23 orthologs with a total of 53 primer pairs for all four species (Table S2, Supporting information) for the first run with a reference. The input files per species (i.e. putative orthologous sequences) ranged from 1,445 to 1,523. We detected 38 markers that covered three of the species (99 primer pairs), 81 markers covering two species (258 primer pairs), and 118 markers that covered any single species (684 primer pairs). For the individual species, *Baetis* sp. had the most markers available (209) of the single- and multi-species markers. There were 136 markers for *Eurylophella* sp., 104 markers for *I. bicolor*, and 87 markers for *E. danica*. The lengths for all markers covering all four species varied between 207 and 997 bp with median of 517 bp, containing between 39 and 298 SNPs per marker with a SNP every four bp on average. Marker length and number of SNPs were correlated with a Pearson's correlation coefficient of 0.96 (Pearson's product-moment correlation *P* < 0.001). Without a reference, DISCOMARK identified 25 orthologs with 66 primer pairs for all four species. Run time for this data set on a Linux client (quad-core Intel i5, 8 GB RAM) was < 50 min.

## Discussion

DISCOMARK is the first stand-alone program of which we are aware that discovers putative single-copy nDNA markers and designs primer pairs based on multiple sequence alignments on a genome-wide scale. The visual output gives guidance on the suitability of each marker i.e. variability within and between species measured as number of SNPs, and information about the included species of each marker. Using this approach, primers can be specifically chosen to match the ‘phylogenetic resolution’, i.e. many markers with intermediate number of SNPs for closely related species, and fewer markers with generally higher number of SNPs for distantly related species can be selected. The automatic processing, including combining, aligning, trimming and blasting sequences of any nucleotide FASTA sequences together with the produced graphical output significantly facilitate the design of primer pairs for a large number of nDNA markers. Nevertheless, users retain a high degree of flexibility by the stepwise nature of the workflow. DISCOMARK is free, open-source software to assist the development of markers for non-model species on the genome scale. We demonstrated the efficacy of our approach for closely related species as well as for members of divergent families within an order of insects. Using a reference genome enabled resolution of intron-exon boundaries but is not a strict requirement for marker design. We strongly recommend carefully checking the selected primer pairs in the alignment viewer. The performance of DISCOMARK will largely depend on the properties of your input data (i.e. fidelity of ortholog prediction, genome complexity, divergence of species.)

### Marker development within the order Ephemeroptera

The usage of DISCOMARK added an extensive set of new potential nDNA markers to the ones that have been used to date for mayfly (Ephemeroptera) phylogenies based on few individual genes (histone 3, elongation factor 1 alpha, phosphenolpyruvate carboxykinase (Vuataz *et al.* 2011; Pereira-da-Conceicoa *et al.* 2012; Vuataz *et al.* 2013)). Most recent tree reconstructions remain based on mitochondrial DNA markers (e.g. Rutschmann *et al.* 2014 (used three mitochondrial genes); Leys *et al.* 2016 (used cytochrome *c* oxidase subunit 1 *(cox1)* gene)). With DISCOMARK, the availability of more genome data will increase the number of markers suitable for phylogenetic studies. This will promote more fine-scale phylogenetic studies, which are needed to resolve more recent evolutionary events and the phylogenetic relationships of morphologically cryptic species that can not be resolved with standard markers (Dijkstra *et al.* 2014).

## Acknowledgements

We thank Katrin Preuß and Susan Mbedi for their help with genome sequencing, and to our research groups, in particular to David Posada and Sara Rocha, and to Eric Coissac and three anonymous reviewers for constructive comments that improved the quality of this project. Research was partially supported by the Leibniz Association (PAKT für Forschung und Innovation “FREDIE” project), the Swiss National Science Foundation (Early PostDoc.Mobility grant P2SKP3_15869 to S.R.), and a visiting fellowship from the Japan Association for the Advancement of Science (L-15543 to M.T.M.). This is publication number 39 of the Berlin Center for Genomics in Biodiversity Research.

## Data Accessibility

The program, user manual and example data sets are freely available at: https://github.com/hdetering/DISCOMARK (last accessed March 28, 2016). Scripts used for the analyses are available at: https://github.com/srutschmann/python scripts (last accessed March 28, 2016). All DNA sequences from this study are available under GenBank accessions: KU987258-KU987260, KU987265-KU987268, KU987273-KU987276, KU987285-KU987288. GenBank accession numbers for sequences included in previous studies are the following:KU971838-KU971840,KU971851, KU972090-KU972092, KU972104, KU972490-KU972492, KU972503, KU973060-KU973061, KU973074. Sequence alignments and tree files are available at the Dryad repository (doi.org/10.5061/dryad.9sf96).

## Author Contributions

S.R., H.D., and M.T.M. conceived the study. S.R. coordinated the project and performed the empirical work. S.R. and H.D. designed the program. H.D. implemented the program. S.R., H.D., and M.T.M wrote the manuscript. S.S. gave guidance for the ortholog prediction. J.F. provided the code of the PRIFIweb tool. All authors provided comments and approved the final manuscript.

